# Decomap-seq enables efficient and reliable retrieval of spatial transcripts

**DOI:** 10.64898/2026.03.18.712548

**Authors:** Kaiqiang Ye, Handong Wang, Wenjia Wang, Xiangwei Zhao

## Abstract

Spatial transcriptomics (ST) has emerged as a transformative tool for resolving the molecular heterogeneity of complex tissues within their native anatomical context. However, next-generation sequencing (NGS)-based ST platforms frequently encounter sensitivity bottlenecks arising from sub-optimal probe architectures on solid substrates. Conventional single-stranded DNA coupling methods often lead to disordered interfacial molecular conformations due to non-specific nucleobase-mediated surface tethering, which creates steric hindrance and inhibits the enzymatic efficiency of in situ library preparation. Here, we present Decomap (double-strand protected combinatorial barcoding microarray chip), a high-performance ST platform utilizing a triple-segment (dsZ-X-Y) fabrication strategy to achieve superior transcript capture efficiency. This structural optimization significantly enhances DNA ligation kinetics and subsequent polymerase-mediated extension, overcoming the fundamental limitations of traditional single-stranded coupling strategies. Decomap-seq achieved a median detection of 7,200 genes and 29,097 UMIs per 50 μm-spot at a sequencing saturation of 50.1%. These results validate Decomap as a highly sensitive and robust tool for spatial transcriptomics, offering a powerful platform for advancing research in histopathology, developmental biology, and neuroscience.

## 1. Introduction

Spatial transcriptomics (ST) has revolutionized biological research by mapping gene expression within its native tissue context, overcoming the inherent loss of spatial information characteristic of traditional transcriptomics^1, 2^. This spatial resolution provides a transformative lens through which to investigate tissue heterogeneity, intercellular signaling, and the spatiotemporal dynamics of disease progression^3-5^. Among ST methodologies, Next-Generation Sequencing (NGS)-based approaches utilizing in situ oligo-dT capture have gained prominence due to their capacity for unbiased, whole-transcriptome profiling.

However, conventional solid-phase synthesis of capture microarrays often relies on single-stranded oligonucleotide coupling. In this “ssX-Y” configuration, amino groups on nucleobases frequently trigger non-specific surface interactions, causing probes to tether in non-terminal orientations^6, 7^. This sub-optimal interfacial conformation not only reduces the ligation efficiency of subsequent combinatorial barcoding probes but also creates steric hindrance during library preparation, inhibiting enzymatic DNA extension and ultimately compromising transcript capture sensitivities.

To address these challenges, we developed Decomap (double-strand protected combinatorial barcoding microarray chip), a platform utilizing a triple-segment (dsZ-X-Y) fabrication strategy. By incorporating a double-stranded regional Z-barcode (dsZ), we effectively shielded the probes from non-specific surface binding during coupling, thereby preserving the sequence integrity and accessibility of the PCR adapters. Furthermore, by re-engineering the microfluidic circuitry and regional barcoding logic, we achieved high-throughput parallel fabrication of multiple barcoded arrays on a single slide, significantly enhancing production efficiency^8^. Benchmarking experiments demonstrated that Decomap significantly outperforms the conventional ssX-Y method in gene detection sensitivity.

In spatial profiling of the mouse brain, Decomap-seq achieved a median detection of 7,200 genes and 29,097 UMIs per 50 μm-spot at a sequencing saturation of 50.1%. These results validate Decomap as a highly sensitive and robust tool for spatial transcriptomics, offering a powerful platform for advancing research in histopathology, developmental biology, and neuroscience.

## 2. Results

### 2.1 Design and fabrication of Decomap

The modification of combinatorial barcoding DNA microarray chip for spatial mRNA sequencing typically involves the covalent coupling of single-stranded barcodes X (Figure 1A) to a functionalized slide with the assistance of microfluidics, followed by the orthogonal ligation of barcodes Y to X (ssX-Y method). However, a distinctive feature of Decomap lies in the covalent of a double-stranded barcodes Z to the slide, followed by two rounds of DNA ligation to orthogonally assemble barcodes X and Y along the x and y axes, ultimately forming the final DNA microarray chip (Fig. 1B). Specifically, in contrast to the ssX-Y method, Decomap first introduces double-stranded barcodes Z regionally onto the slide through covalent coupling.

**Figure 1.**
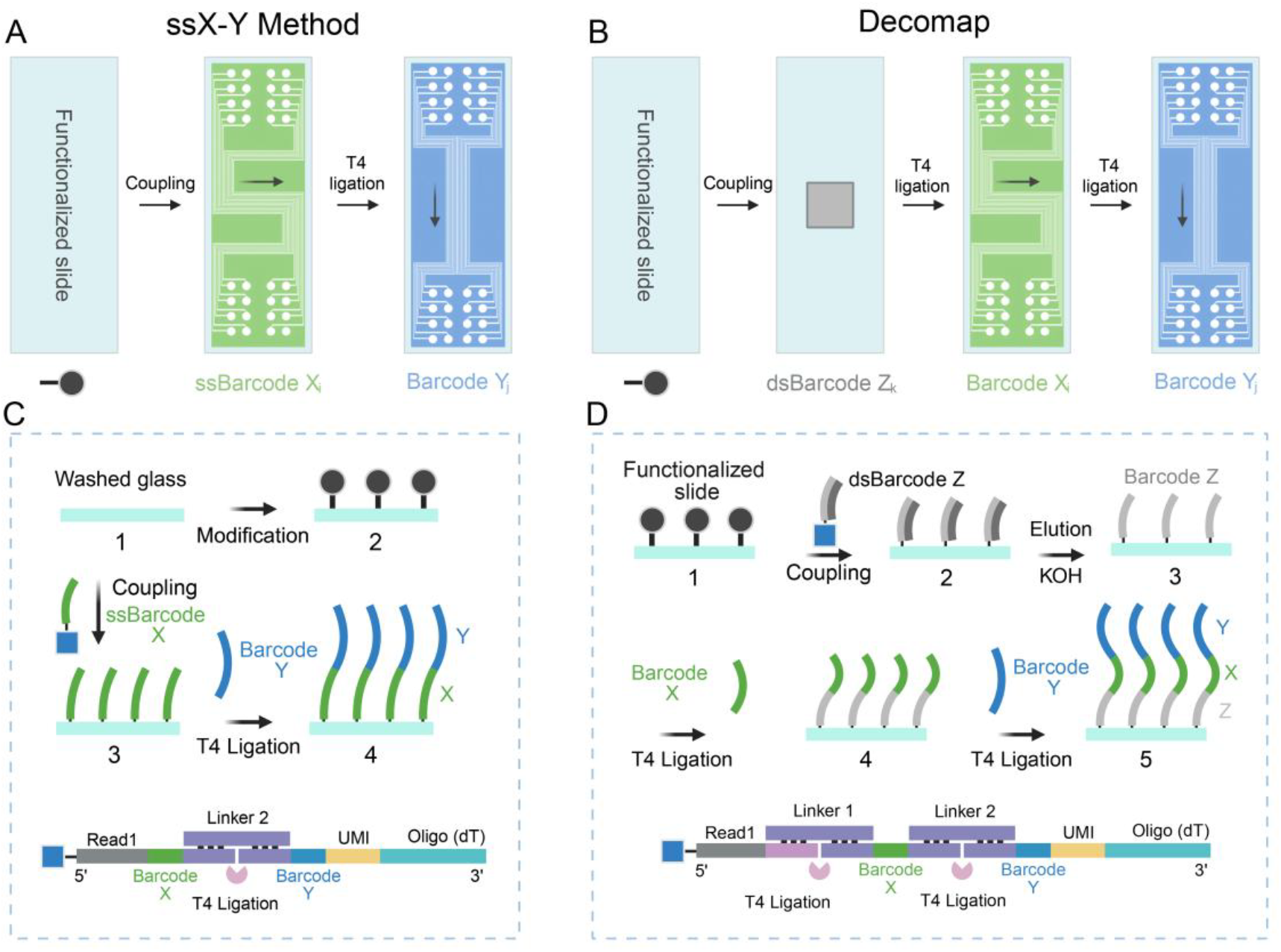
Workflows and details of ssX-Y method and Decomap.

Subsequently, KOH is used to denature and elute the complementary probes, thereby retaining the intact read1 sequences that are coupled via the 5’ terminal groups on the substrate. A series of barcodes X and Y are then guided by PDMS channels to the barcodes Z-modified regions, where two sequential T4 DNA ligation reactions enable in situ combinatorial assembly along the x and y axes, resulting in a three-fragment combinatorial barcoding microarray chip. Additionally, barcodes Z act as regional labels, facilitating simultaneous preparation of multiple combinatorial barcoding microarrays on one single slide. This approach not only enhances chip fabrication efficiency and modification uniformity but also enables concurrent construction of sequencing libraries from different samples, minimizing experimental deviations and reducing library preparation costs.

### 2.2 Decomap enhances the retrieval of spatial mRNA

Our fabrication workflow begins with the functionalization of aminated substrates into double-stranded anchoring linkers (dsBarcode Z), engineered for high coupling efficiency and minimal non-specific adsorption. Subsequently, high-throughput DNA-barcoded microarrays were synthesized via two rounds of microfluidic-guided orthogonal DNA ligation (Fig. 2A). The CAD schematics, PDMS microfluidic hardware, and reaction assembly are detailed in Fig.2 B-C. Fluorescence in situ hybridization verified exceptional chip uniformity with negligible background noise (Fig. 2D). Quantitative fluorescence analysis (Fig. 1E) further confirmed consistent modification across distinct array regions (n = 10), underscoring the high reproducibility and stability of the Decomap fabrication process. To verify the efficacy of the double-strand protection strategy in preventing non-specific coupling of DNA probes, we compared the reaction results for three types of DNA probes immobilized on a substrate via DSS (Fig. 2F). These probes included single-stranded read1, double-stranded read1 with 5’ amino modification, and unmodified double-stranded read1. Following probe immobilization, elution was supporting our hypothesis. In contrast, the unmodified double-stranded read1 showed no significant signal, indicating that the amino groups on the bases did not react in the double-stranded form. These results demonstrate that, in the immobilization process of single-stranded barcodes X in the ssX-Y method, there are many situations in which the amino sites on the bases were immobilized on the substrate, while the double-strand protection strategy effectively avoids non-specific probe coupling. Benchmarking experiments in mouse brain tissue demonstrated that the dsZ-X-Y strategy yields a marked improvement in transcriptomic capture sensitivity compared to the ssX-Y method (Figs. 1G). This performance gain is directly attributed to the reduction of non-specific coupling sites and optimized DNA ligation kinetics. Notably, the fabrication and operational costs of this platform are substantially lower than those of existing commercial spatial transcriptomics solutions, offering a cost-effective and high-performance tool for large-scale spatial molecular profiling.

**Figure 2.**
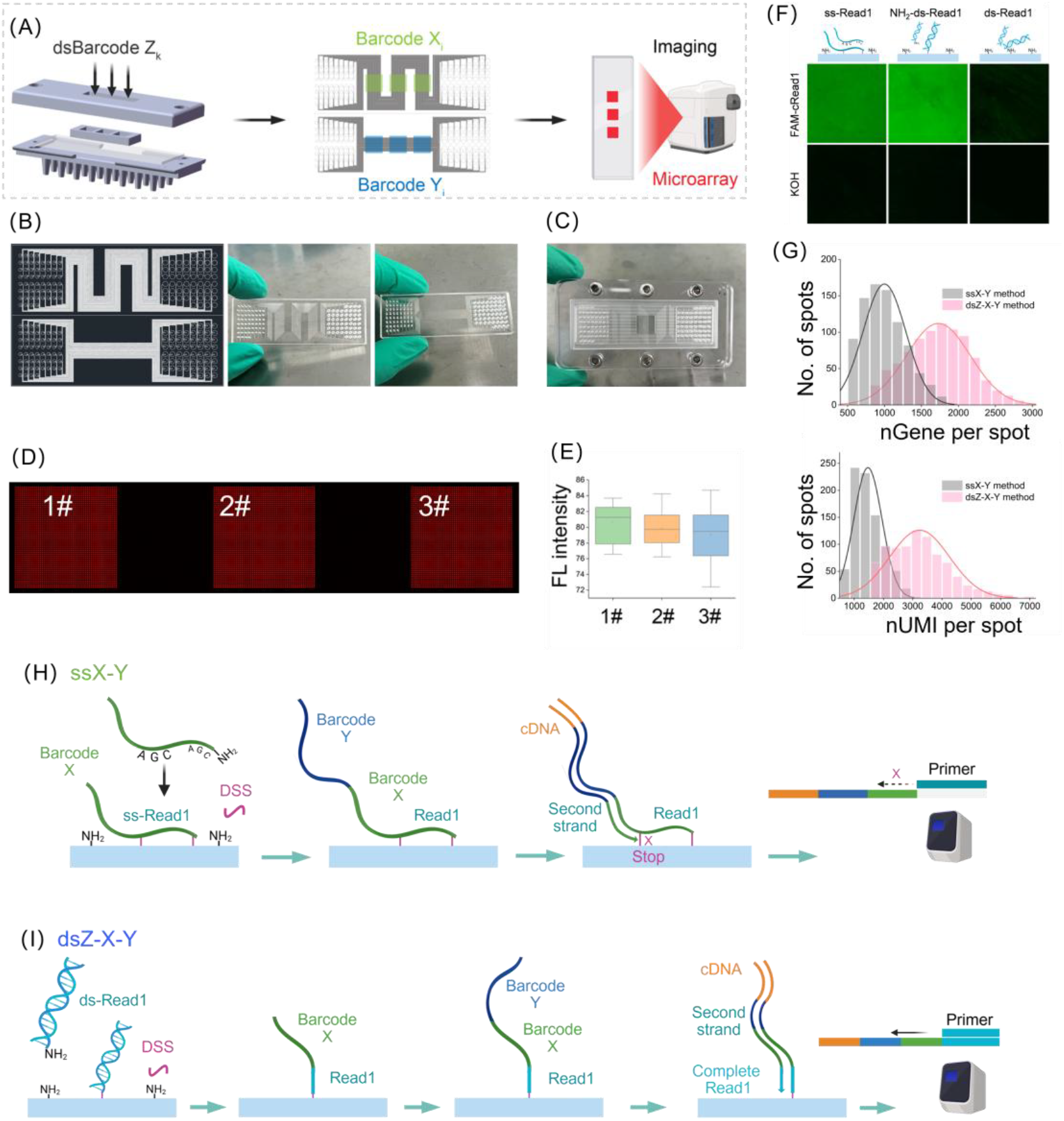
Fabrication workflow and performance characterization of the Decomap platform. (A) Schematic illustration of the spatial transcriptomics (ST) chip fabrication process using the dsZ-X-Y strategy. (B) CAD schematic (left) and representative photograph (right) of the PDMS microfluidic device. (C) Photograph of the customized reaction fixture for in-chip DNA ligation. (D) Full-slide fluorescence scan of the ST chip following hybridization with Cy3-labeled PolyA probes. (E) Quantitative analysis of fluorescence intensity across distinct array regions (n = 10 independent areas), demonstrating high spatial uniformity. (F) Fluorescence images of different probe configurations (ss-Read1, NH_2_-dsRead1, and ds-Read1) after covalent coupling to aminated substrates, hybridization with FAM-cRead1 probes, and KOH stringency washing. (G) Comparison of transcriptomic capture efficiency between the dsZ-X-Y and ssX-Y fabrication methods. (H, I) Comparative analysis of the dsZ-X-Y and ss-X-Y strategies; the double-stranded anchoring linker minimizes non-specific coupling sites, significantly enhancing the transcriptomic recovery performance of the ST chip.

### 2.3 Decomap-seq obtains precise spatial gene expression patterns in tissue sections

Spatially Resolved Transcriptomic Profiling of the Mouse Brain To evaluate the biological fidelity of the platform, we performed unsupervised clustering and spatial mapping of the gene expression data. The resulting spatial clusters in the mouse brain and olfactory bulb exhibited high concordance with the anatomical annotations in the Allen Brain Atlas, validating that dsZ-X-Y-seq faithfully recovers complex gene expression patterns (Figs. 3C-E). The spatial distributions of representative heterogeneity marker genes are shown in Figs. 3F-G. Notably, high-resolution dsZ-X-Y-seq (15 μm-spots) enabled the precise identification of hippocampal sub-structures and the spatially resolved analysis of cell-type-specific markers (Figs. 3H–I). Collectively, these results demonstrate that our unique modification and barcoding strategy endows dsZ-X-Y-seq with superior detection sensitivity, facilitating high-throughput spatial transcriptomic mapping at near-single-cell resolution.

**Figure 3.**
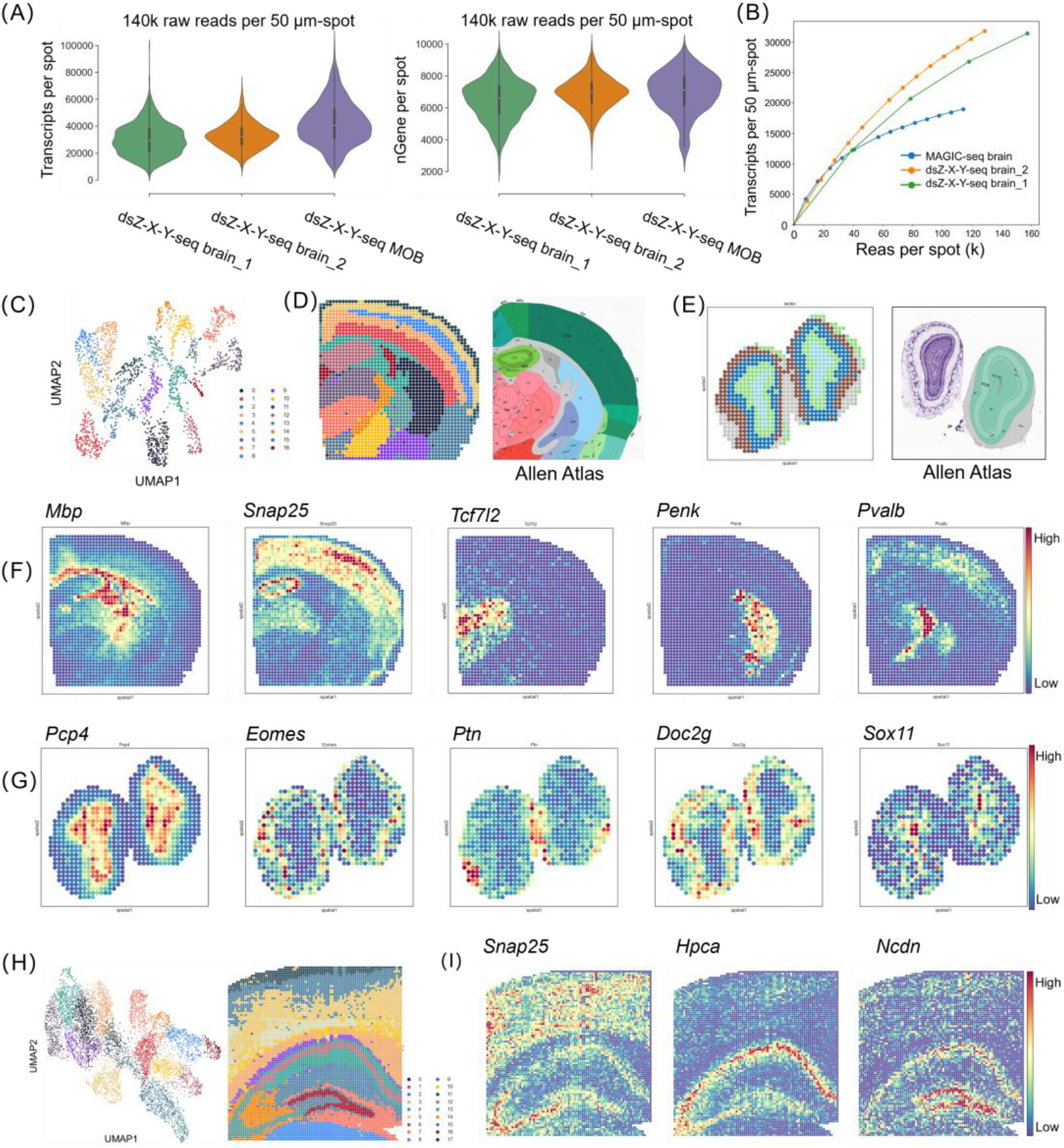
High-performance gene detection and spatial transcriptomic mapping via dsZ-X-Y-seq. (A) Gene detection sensitivity of dsZ-X-Y-seq in mouse brain and olfactory bulb sections (standardized at 140k raw reads per 50 μm-spot). (B) Rarefaction curves demonstrating the robust transcriptomic recovery of dsZ-X-Y-seq across increasing sequencing depths. (C, D) Unsupervised clustering and spatial projection of gene expression profiles in the mouse brain, showing high anatomical concordance with the Allen Brain Atlas. (E) Spatial clustering and visualization of the mouse olfactory bulb validated against corresponding anatomical regions in the Allen Brain Atlas. (F, G) Spatial expression patterns of representative marker genes exhibiting regional heterogeneity in the mouse brain and olfactory bulb. (H, I) High-resolution spatial profiling (15 μm-spot) of the mouse hippocampal formation, enabling the precise deconvolution of sub-structural organizations and marker gene distributions.

## 3. Discussion

The prominent advantage of Decomap lies in its remarkable transcript retrieving in spatial mRNA sequencing. In the case of disuccinimidyl suberate (DSS)-coupled modification (Fig. 2H), the ssX-Y method immobilizes the read1 (a part of the barcodes X) onto the substrate through DSS-mediated crosslinking between the amino groups of both the substrate and the barcodes. However, among the four nucleotide bases (A, T, C, and G), all except thymine (T) possess primary amines capable of reacting with DSS. Thus, it is possible that the barcodes X is crosslinked to the substrate through amino groups in the base rather than the 5’-terminal modified amino group, or that both the 5’-terminal amino group and base-derived amino groups are simultaneously reacted with DSS and crosslinked to the substrate. In such cases, as long as the linker at the 3’ end of the barcodes X remains functional, it allows to continue to ligating with the barcodes Y, which can capture mRNA molecules in subsequent sequencing experiments. However, during the amplification reaction, cDNA generated from such non-specifically crosslinked probes may partially fail to amplify due to the lack of primer-binding sites, ultimately leading to the loss of captured transcripts. In the Decomap-seq, the amino groups on the bases are protected by the complementary strand of the double-stranded barcodes Z (Fig. 2I), ensuring that only the 5’-terminal modified amino group participates in the reaction during DSS crosslinking. These specifically crosslinked probes ensure the successful amplification of cDNA generated from the captured transcripts in spatial RNA sequencing, thereby increasing the complexity of the sequencing library and enhancing gene detection efficiency.

## 4. Methods

### 4.1 Preparation of amino-modified slides

The slides were thoroughly washed, dried, and then placed into a glass staining tank. Piranha solution was prepared in a fume hood by mixing concentrated sulfuric acid and 30% hydrogen peroxide in a 7:3 volume ratio. Sulfuric acid was added first, followed by the gradual addition of hydrogen peroxide. The prepared solution was then transferred into the staining tank, where the slides were fully submerged and left to react for 2 hours. Following the treatment, the slides were rinsed extensively with ultrapure water. The dried slides were subsequently immersed in a 2% (v/v) APTES ethanol solution and allowed to react at room temperature for 3 hours. Afterward, the slides were rinsed with absolute ethanol, subjected to ultrasonic cleaning for 3 min, washed thoroughly with ultrapure water, and dried under a stream of nitrogen. The vacuum drying oven was preheated to 110°C, and the dried slides were placed in the oven to bake for 1 hour. Following natural cooling, the slides with amino-modified surfaces were obtained and could either be stored at 4°C or directly used for subsequent processing.

### 4.2 Design and fabrication of the microfluidic devices

The microfluidic devices were fabricated based on soft lithography and polydimethylsiloxane (PDMS) casting techniques. Briefly, the layout for X-channel along the x-axis and Y-channel along the y-axis of the slide were designed using AutoCAD. The resulting blueprint was outsourced to Suzhou Yancai Micro-Nano Technology Co., Ltd. for the fabrication of a silicon wafer template. Upon conversion of the layout patterns into corresponding photomasks, the outer channel regions were subjected to wet etching on 4-inch silicon wafers through a series of processes, including photoresist spin-coating, lithography, and development. The pattern on the silicon template served as mold for subsequent fabrication of the PDMS microfluidic channels.

Approximately 35 ml of PDMS and its curing agent, mixed in a 10:1 mass ratio, were combined thoroughly in a plastic cup and poured onto the silicon template. The mixture was allowed to stand for 5 min to ensure complete spreading across the silicon surface. The mold was then transferred to a vacuum oven for 10 min of degassing, followed by incubation at 4°C for 10 min to eliminate air bubbles on the PDMS surface. The oven was preheated to 68°C, and the PDMS solution was cured for 2 hours, after which the sample was removed and allowed to cool naturally. Using a cutter knife, the PDMS was cut along the outermost contour lines, resulting in two separate pieces for the X and Y microfluidic channels. The PDMS was placed on a hole puncher, and holes were created using 1.0-mm or 1.5-mm diameter punch. Any surface impurities were removed with high-pressure gas before the PDMS was stored in a clean tip box. Additionally, the polycarbonate (PC) and aluminum alloy components for the probe modification reactions in the PDMS channels, as well as the subsequent ST reaction, were designed in SolidWorks and custom-fabricated using CNC (computer numerical control) machining.

### 4.3 Generation of Decomap with dsZ-X-Y method

To obtain the double-strand barcode Z (dsZ), the amino-modified single-strand barcode Z and its completely complementary probe (CZ) were dissolved and subjected to annealing in a thermocycler with a molar ratio of 1:1. The dsZ conjugation solution, which inclued 30 μM dsZ solution, 1×PBS (pH 7.0) and 2.5 mM DSS solution, were added to the reaction chambers and incubated at room temperature for 1 hour. The slide was washed three times with buffer 1 (2× SSC containing 0.1% SDS), followed by rinsing with ultrapure water and drying under a stream of nitrogen. The slides were blocked in 10% (w/v) succinic anhydride solution (containing 1% v/v triethylamine) at room temperature for 45 min, then washed with ultrapure water and dried with nitrogen. The dsZ-modified regions were denatured with 0.08 M KOH for 2 min, followed by washing with 0.1× SSC and ultrapure water. The modification regions were marked using a diamond pen for the following image alignment.

Then, barcode X solutions (X1-50) were annealed with Linker A to obtain XL solutions, which were dispensed into 96-well plates for storing. The X microfluidic channels were aligned and attached to the dsZ-modified slide, and the images of the X channels and marks were acquired using a microscope (Olympus, IX73) in bright field. The ligation solutions, which consist of 20 μM of XL solution, 1× T4 ligation buffer and 40 U μl^−1^ T4 DNA ligase, were added to the inlet holes and aspirated into the microfluidic channels. The ligation reaction was performed in a humid chamber at 37°C for 30 min or room temperature for 2 h, followed by washing with buffer 1 and ultrapure water.

To generate the arrays, The Y microfluidic channels perpendicular to the direction of barcode X were aligned and attached to the barcode X-modified slide, and the images of the Y channels and marks were acquired. The ligation solutions, which consist of 10 μM of YL solution (annealed with Linker B), 1× T4 ligation buffer and 40 U μl^−1^ T4 DNA ligase, were added to the inlet holes and aspirated into the microfluidic channels. The subsequent treatments were identical to the steps in the barcode X reaction. The barcoding regions were denatured with 0.08 M KOH for 30 s to remove unligated probes. The washed and dried chip was then placed in a sealed package with a desiccant and stored at 4°C.

### 4.4 Animal experiments

The animal treatments in this study were approved by the Animal Ethics and Welfare Committee of Zhongda Hospital, Southeast University (20200104005) and performed in strict accordance with the guidelines. All applicable institutional and national regulations on the care and use of animals were adhered to. Male mice (6-10 weeks old, C57BL/6J) were purchased from Nanjing Qinglongshan Animal Farm. Before the experiment, mice were bred at 25°C with ad libitum access to food and water to acclimate to the environment. Mice were euthanized by cervical dislocation, brain and MOB were immediately isolated and embedded in optimal cutting temperature (OCT) compound, snap-frozen with liquid nitrogen and then stored at -80°C.

### 4.5 Tissue sectioning and H&E staining

For sectioning, the tissue block was removed from the -80°C freezer and placed on the cold stage of a cryostat. The temperatures of the cold stage, sample head and blade were set at -25°C, -11°C and -18°C, respectively. The tissue block was adhered to the sample chuck and equilibrated to the sample head temperature (−11 °C) for 20 min. After trimming the tissue to the target area, serial sections were prepared at a thickness of 10 μm and mounted onto the Decomap.

Sections were incubated on a thermocycler adaptor at 37°C for 1 min, fixed in prechilled methanol at −20 °C for 30 min. Then, the sections were treated with isopropanol for 2 minutes and air-drying for the following staining of hematoxylin for 2 min, bluing buffer for 1min and eosin for 30 s, followed by washing with ultrapure water after each stain. After incubation at 37 °C for 5 min, images of the sections and the marks were acquired by scanning under brightfield.

### 4.6 Tissue permeabilization and RT

To determine the optimal tissue permeabilization time, tissue optimization (TO) experiment was performed. First, the H&E stained tissue sections were treated with 0.1% pepsin (diluted in0.1 M HCl) at 37°C across different time gradients, then Cy3-dCTP-labeled cDNA synthesis was performed by adding 50μl of TO reverse transcription (RT) mix to pepsin-pretreated tissue sections. The TO RT mix contained 10 μl of 5× RT buffer, 2.5 μl of TO dNTP mix (dATP/dGTP/dTTP at 500 mM each, dCTP at 12.5μM), 0.5 μl of GTP (100 mM), 1.5 μl of template switch oligo (100 μM), 2.5 μl of recombinant RNase inhibitor (40 U μl^-1^), 1.25 μl of Cy3-dCTP (1 mM), 2.5 μl of Maxima H Minus RTase (200 U μl^-1^), and 29.25 μl of water. After reaction at 42 °C for 1.5 h, sections were washed three times with 0.1× SSC buffer, followed by addition of 50 μl of proteinase K solution (16 U ml^− 1^) and incubation at 37°C for 30 min. Then, each well was sequentially washed twice with buffer 1, three times with 0.1× SSC, and three times with ultrapure water, then dried under nitrogen stream. Cleaned chips were scanned using a fluorescence microscope (Olympus IX73), the permeabilization time corresponding to images with the strongest fluorescence signal, highest contrast, and minimal signal diffusion was selected as the optimal condition. RT for library construction was performed at the same cDNA synthesis condition except replacing the TO dNTP mix and Cy3-dCTP with 2.5 μl of dNTP mix (10 mM each).

### 4.7 Secondary strand synthesis

After cDNA synthesis, the wells were washed twice with 0.1× SSC, followed by adding 50 μl of KOH (0.08 M) and incubation for 5 min to denature the mRNA/cDNA strands. After washing three times with 0.1× SSC, the second-strand synthesis mix was added, which contained 5 μl of 10× buffer, 5 μl of dNTP mix (10 mM), 1.25 μl of second-strand primer (100 μM), 2 μl of MgSO_4_ (100 mM), 34.75 μl of water, and 2 μl of Bst 2.0 polymerase. The reaction was carried out at 65°C for 15 min. To collect the second-strand cDNA, followed by three washes with 0.1× SSC, 40 μl of KOH (0.08 M) was added and incubated for 10 min. The KOH was pipetted up and down 10 times within the reaction chamber, then transferred to a 200-μl tube and neutralized with 6 μl of tris buffer (1 M, pH 7.0).

### 4.8 Library construction and sequencing

Subsequently, 50 μl of 2× Kapa HotStart HiFi ReadyMix and 2 μl each of primer (20 μM) were added to the tube above. The mix was subjected to PCR amplification using the following program: 95°C for 3 min; 11-14 cycles of 98°C for 20 s, 65°C for 30 s, and 72°C for 2 min, followed by a final extension at 72°C for 5 min and holding at 4°C. After reaction, 60 μl (0.6×) of VAHTS DNA Clean Beads were added to each tube to purify the PCR products, followed by the final air-drying, 20.5 μl of water was added to elute the DNA products. The concentration and fragment analysis were performed using Qubit dsDNA assay kit and Fragment Analyzer, respectively.

To generate the sequencing libraries, 5 ng of DNA product was processed using a TruePrep Homo-N7 DNA Library Prep kit according to the manufacturer’s protocol. The generated libraries were then sequenced on an Illumina NovaSeq 6000 platform in paired-end 150-bp (PE150) mode.

